# The G-box transcriptional regulatory code in *Arabidopsis*

**DOI:** 10.1101/128371

**Authors:** Daphne Ezer, Samuel JK Shepherd, Anna Brestovitsky, Patrick Dickinson, Sandra Cortijo, Varodom Charoensawan, Mathew S. Box, Surojit Biswas, Philip Wigge

**Affiliations:** Sainsbury Laboratory, University of Cambridge, 47 Bateman St., Cambridge CB2 1LR, UK.; Department of Plant Sciences, University of Cambridge, Downing St., Cambridge CB2 3EA, UK.; Department of Biochemistry, Faculty of Science, and Integrative Computational BioScience (ICBS) center, Mahidol University, Bangkok 10400, Thailand.

## Abstract

Plants have significantly more transcription factor (TF) families than animals and fungi, and plant TF families tend to contain more genes—these expansions are linked to adaptation to environmental stressors (1, 2). Many TF family members bind to similar or identical sequence motifs, such as G-boxes (CACGTG), so it is difficult to predict regulatory relationships. We determine that the flanking sequences near G-boxes help determine *in vitro* specificity, but that this is insufficient to predict the transcription pattern of genes near G-boxes. Therefore, we construct a gene regulatory network that identifies the set of bZIPs and bHLHs that are most predictive of the gene expression of genes downstream of perfect G-boxes. This network accurately predicts transcriptional patterns and reconstructs known regulatory subnetworks. Finally, we present Ara-BOX-cis (araboxcis.org), a website that provides interactive visualisations of the G-box regulatory network, a useful resource for generating predictions for gene regulatory relations.

## INTRODUCTION

Many transcription factors (TFs) are part of large families, with many members binding to highly overlapping sets of binding sites. Therefore, any change in a TF’s concentration or its spatial or temporal distribution may result in unexpected cross talk within the gene regulatory network—a TF may inadvertently affect the gene expression of gene targets of its other family members. This cross-talk phenomenon within TF families appears universal within the eukaryotes, and has been described in yeast (3), plants (4) and mammalian cancer cell lines (5). Understanding the mechanisms that govern how TFs within large TF families regulate their target genes is therefore an important challenge.

Understanding gene regulatory network in plants is further complicated by the fact that plants have more and larger TF families than animals or fungi (1), and even larger families than would be expected through whole genome duplication alone (1, 6). For instance, the highly conserved G-box motif (CACGTG) is bound by TFs in the basic-helix-loop-helix (bHLH) and basic leucine zipper (bZIP) families in organisms ranging from yeasts to humans. However, in plants these two TF families have massively expanded—for example, the bHLH family is now the second largest TF family in plants, with over 100 members in *Arabidopsis*, despite having arisen from an estimated 14 founder genes in ancient land plants (7). At least 80 of these bHLHs have the precise amino acid composition in their DNA-binding domain required to bind to G-box elements (7, 8). Many of the other bHLHs may bind to E-box elements, which retain four nucleotides of the G-box core (ACGT or CANNTG). The bZIP family has similarly expanded from 4 founder genes to over 70 (9).

Moreover, both bHLHs and bZIPs bind to DNA as either homodimers or heterodimers—further increasing the possible regulatory combinations. Even non-G-box binding bHLHs and other HLHs can indirectly regulate G-box-regulated genes, by competing with G-box binding bHLHs for dimerisation partners (10, 11). Furthermore, bZIPs and bHLHs can act antagonistically, competing for binding to the same sites—such as the competition for binding to targets shared between the bHLH PHYTOCHROME INTERACTING FACTOR 3 and the bZIP ELONGATED HYPOCOTYL 5 (12).

Understanding how bHLHs and bZIPs regulate their downstream target is critical, as these TFs have been identified as key regulators of growth (13), temperature response (14), immune response (15), metal homeostasis (16), drought response (17), and light signalling (14, 18)—pathways that would serve as promising targets for improving crop yield and environmental resilience. Much of our knowledge of G-box specificity comes from studies in yeast and mammalian systems, and these results suggest that bZIPs and bHLHs are affected by the local shape of the DNA flanking the binding site (3, 19) and epigenetics (20, 21). Since these families are much larger in plants than in the other organisms studied, it is unclear whether these mechanisms are sufficient to explain TF-to-binding site interactions in *Arabidopsis*. This has been a hindrance to the plant research community, as there have been many cases where a G-box has been identified as critical for the regulation of a gene of interest, but researchers were unable to distinguish between the many possible candidate TFs that might regulate that gene (22–26).

In this paper, we identify a set of approximately 2000 genes that are highly likely to be regulated by an upstream G-box motif in *Arabidopsis* seedlings, and we ask how bHLHs and bZIPs might regulate these genes. We find that while the sequences immediately flanking the G-boxes enable us to predict the *in vitro* binding of bZIP homodimers, the flanking sequences are not sufficient to explain the gene expression profiles of the downstream genes. Therefore, we constructed a network to identify the bHLHs and bZIPs whose expression was most predictive of the expression profiles of genes downstream of perfect G-box motifs. This network can predict a number of well-established sub-networks, and the entire network can be browsed through an interactive network visualisation website we have developed, called Ara-BOX-cis. This study provides useful resources for those who find a G-box near their gene or genes of interest without clear bHLH or bZIP targets, which is often a roadblock to understanding the regulation of their genes. By utilising these novel tools, we can help researchers make testable predictions of gene regulation interactions.

## RESULTS

### bZIP homodimers only bind to a subset of perfect G-boxes *in vitro*

We sought to identify a comprehensive set of genes likely to be regulated by G-box binding TFs in *Arabidopsis* seedlings as our reference set. For this, we selected those genes containing a perfect G-box sequence (CACGTG) within 500bp upstream of their transcription start site (TSS) and that were expressed in a substantial number of samples of 7-day old seedlings at 22^°^C (see “selection of genes” section in Methods and **Fig S1-A**). A total of 2146 genes fit these criteria **(Table S1)**. Although there are many known cases where TFs affect targets more than 500bp away from their binding sites, TFs that bind to proximal sites are more likely to affect the transcription of the nearby gene. Moreover, there is enrichment for G-box sequences within 500bp of a TSS (**Fig S1-B**).

Firstly we sought to determine whether bHLHs and bZIPs are capable of binding to all perfect G-boxes, or only a subset of these sites. To resolve this, we compared *in vitro* binding of bZIPs, bHLHs, and BZR (a TF that is not a bHLH or bZIP but whose binding sites are CGTG, a subset of the G-box (41)). We identified binding sites within our set of promoters containing perfect G-boxes, using previously published DAP-seq data (35) **(Table S2)**. Both methylated and un-methylated DNA sequences were used in this analysis, but methylation had minimal effects on the binding behaviour **(Fig S2)**.

These results indicate that there is a set of bZIPs that only bind to a subset of perfect G-boxes (**Fig 1A**, binding pattern 1), including known strong G-box binders such as G-BOX BINDING FACTOR 3 (GBF3) and bZIP16. This indicates that TFs can distinguish between perfect G-boxes based on the genomic context of the binding site. Fewer bHLH TFs are represented in the DAP-seq data and they often have fewer identified binding events—those that have a fraction of reads in peaks (FRiP) of less than 5 are indicated in orange in **Fig 1A**.

**Figure 1:**
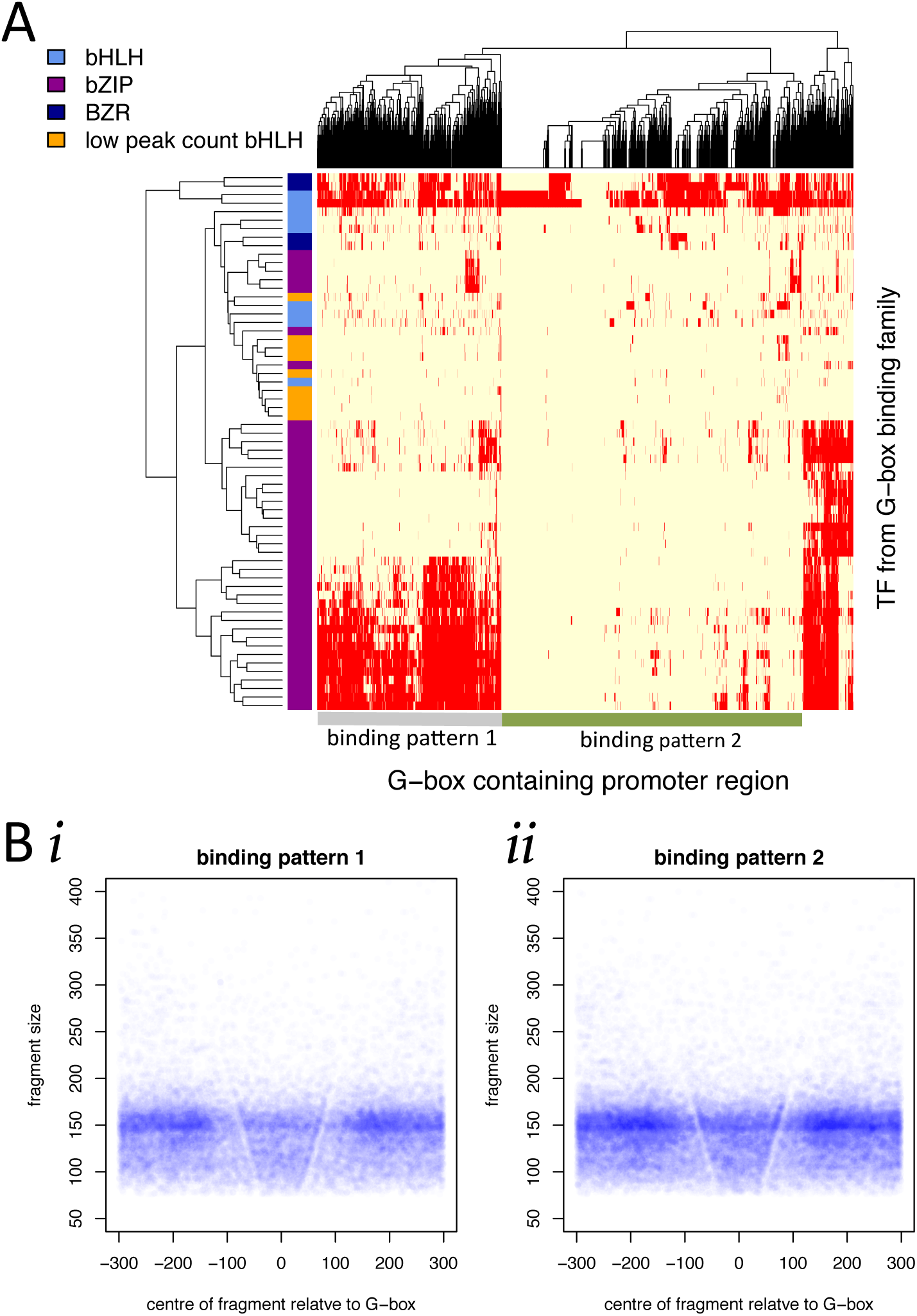
While TFs can differentiate between ‘perfect’ G-box motifs given genomic context, multiple TFs are capable of binding to the same G-box sequence. (A) DAP-seq experiments from (35) illustrate which bZIPs, bHLHs and BZRs are found within 500bp of the TSS of genes with perfect G-boxes in their promoters— a binding event is drawn in red. Note that many bHLH TFs had very low numbers of peaks in the DAP-seq experiment (Fraction of Reads in Peak (FRiP)<5) and these are highlighted in orange. The two largest clusters include binding pattern 1, which has a large amount of bZIP binding and binding pattern 2, which has almost no binding in this DAP-seq experiment. (B) MNase-seq data taken at ZT0 (immediately before dawn) illustrates that there is TF binding *in vivo* in promoters with binding pattern 1 and binding pattern 2, as indicated by the prominent V-shape.

Both BIM2 (a bHLH) and BZR exhibit more ubiquitous binding to G-boxes than the bZIPs. However, there is still a cluster of G-boxes in our set that had very little binding in this dataset (labelled binding pattern 2). Since the DAP-seq data suggests that some G-boxes are not strongly bound *in vitro*, we sought to determine if these sites are likely to be functional *in vivo*, where other factors may act cooperatively to enhance TF binding. Protein-DNA binding information can be inferred from micrococcal nuclease (MNase) data, since proteins that bind DNA protect it and perturb the MNase digestion pattern. In this way, plotting the fragment sizes over genomic position relative to a binding site reveals a characteristic ‘v’ shape if the binding site is occupied (See **Fig S3** for further description of v-plots (42, 43)). For both G-boxes in binding pattern 1 and binding pattern 2, we see a strong ‘v’ pattern suggestive of *in vivo* occupancy (**Fig 1B**). This indicates that G-boxes from both binding patterns may have regulatory roles *in vivo*, but from this data it is not possible to identify the specific TF.

In summary, the DAP-seq data suggests that bZIPs are capable of distinguishing between perfect G-boxes based on genomic context—independent of DNA accessibility or the presence of binding partners. However, many of the G-boxes without strong bZIP binding *in vitro* are still bound by TFs *in vivo*, suggesting other factors might interact with bZIP TFs and influence their binding.

### Sequences flanking G-box can predict whether bZIP homodimers are capable of binding to the sequence

*In vitro* bZIP TFs are able to distinguish between different perfect G-boxes, suggesting additional information is present in the genomic context. We tested whether it is possible to predict bZIP binding based on the sequences flanking the G-box (**Fig 2A**). This has been shown to be a key feature of G-box binding in yeast and mammals (3, 19), and even in plants there are specific examples of this-- e.g. ABRE prefers a GC flank (44). The G-boxes in binding pattern 1 have a slight preference for GA flanks as shown by the depiction of the Position Weight Matrix (PWM) (**Fig 2B**) which is consistent with earlier observations of certain bZIP preferences (45); however, there are many examples of G-boxes without a GA or TC flank that are still bound by these bZIPs. In fact, 48% of G-boxes in binding pattern 1 have neither a ‘GA’ prior to the G-box, nor a ‘TC’ after the G-box, while 16% of G-boxes in binding pattern 2 do have either a ‘GA’ or a ‘TC’ flanking sequence, so this is not sufficient to account for how bZIPs distinguish between G-boxes.

**Figure 2:**
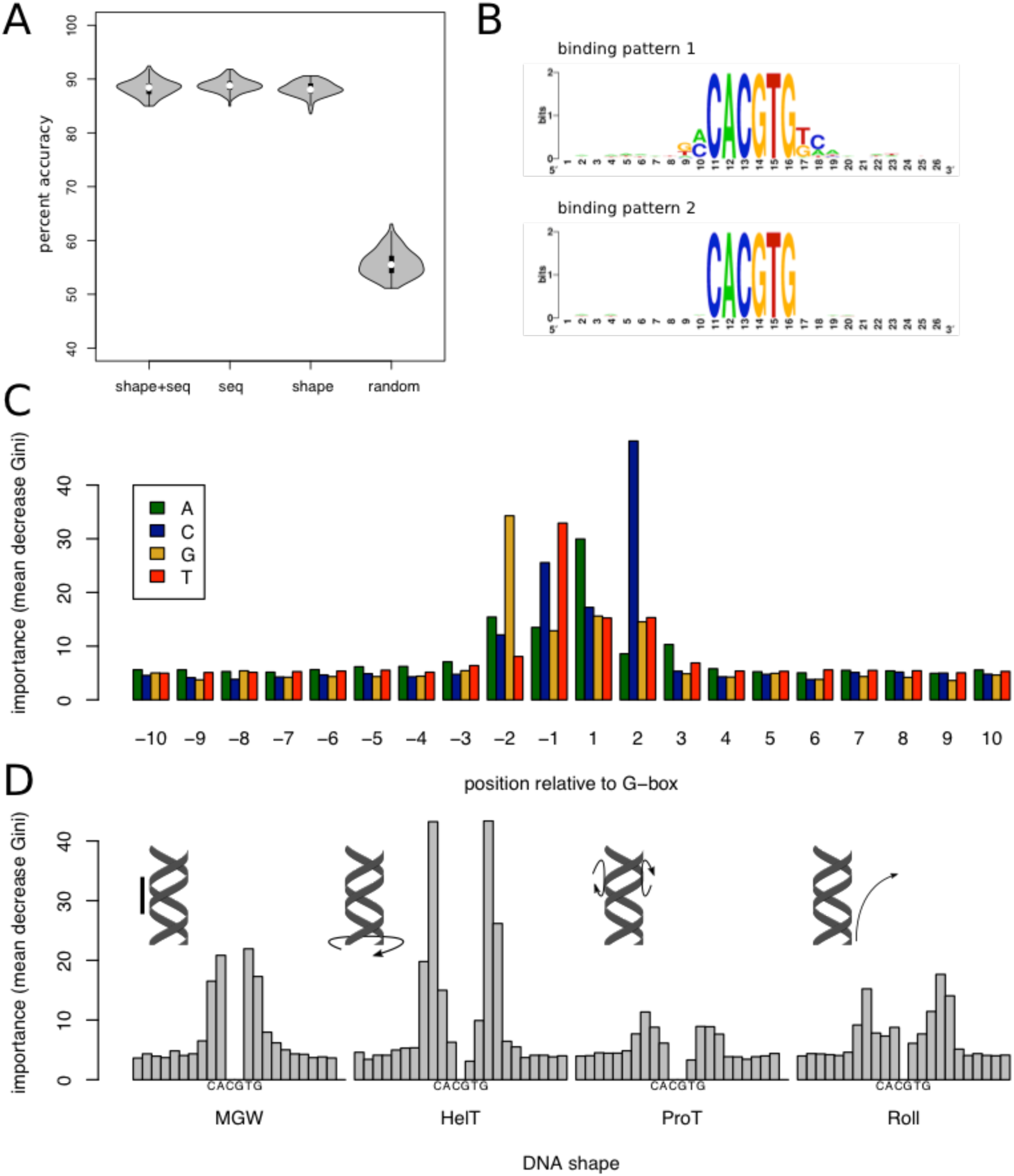
Flanking sequence distinguishes DAP-seq binding patterns with high accuracy. (A) Random forest models using G-box flanking sequence and local DNA shape can distinguish the DAP-seq binding patterns with approximately 88% accuracy. (B) The sequences with many bZIPs bound seem to be enriched in CA and TC flanking sequences (C) The sequence based random forest model suggests that the two bases flanking the G-box are most critical in predicting bZIP binding. (D) The DNA shape features that are most important for distinguishing bZIP bound G-boxes are the minor groove width (MGW) and the helical turn (HelT) pattern.

We then made a predictive model that implicitly handled interaction between bases, using a random forest machine learning approach. Random forests could predict whether or not bZIPs could bind to a particular G-box with an average accuracy of 88% (**Fig 2A**), much higher than would be expected by chance (See **Fig S4** for experimental design). The most informative features in the model were actually the presence of a ‘G’ or a ‘C’ two bases away from the G-box (**Fig 2C**), even though these bases appear less informative than the bases immediately flanking the G-box in the PWM model (**Fig 2B**).

In addition, we considered a model that incorporated shape features such as the width of the DNA sequence or the ability to twist or roll (36)—this performed equally well as the DNA sequence model (**Fig 2A**). We found that helix twist (HelT) and the minor groove width (MGW) were the most important features (**Fig 2D**), as opposed to roll and propeller twist (ProT). In yeast, MGW was found to play a role in determining overall binding affinity of a G-box, but ProT could be used to distinguish between the binding of different G-box binding transcription factors (3). In mammals, the roll feature could be used to predict the binding affinity of Max, a TF that binds to the ACGT core sequence but prefers CACGTG, the complete G-box (19). This suggests that TFs from different species may prefer distinctive DNA shape features.

### Gene expression network predicts bHLH and bZIP regulatory targets

Even though we can predict whether bZIP homodimers are likely to bind to a perfect G-box in the DAP-seq experiment using the flanking DNA sequences, the DAP-seq data does not fully explain how bZIPs and bHLHs regulate genes downstream of G-boxes *in vivo*. Firstly, it is unclear which TFs are binding to the G-boxes in ‘binding pattern 2’ *in vivo*. Secondly, even if a TF is capable of binding to a motif, this does not indicate that this TF is necessarily influencing the gene expression of the target gene. For instance, a TF may bind to a DNA sequence under many conditions, but the TF may only be active/functional under specific conditions. The converse might also be true—a transiently bound TF may still influence the gene expression of a nearby gene via the ‘hit-and-run’ mechanism (46). In our case, we’ve seen that many bZIPs are capable of binding to the same subset of G-boxes, yet it is unlikely all these binding events are biologically relevant. There may be temporal or spatial differences in the concentration of co-factors or in the chromatin near these G-boxes, and these bZIPs have very different gene expression patterns from one another (**Fig 3A**), so they may still regulate different subsets of genes.

**Figure 3:**
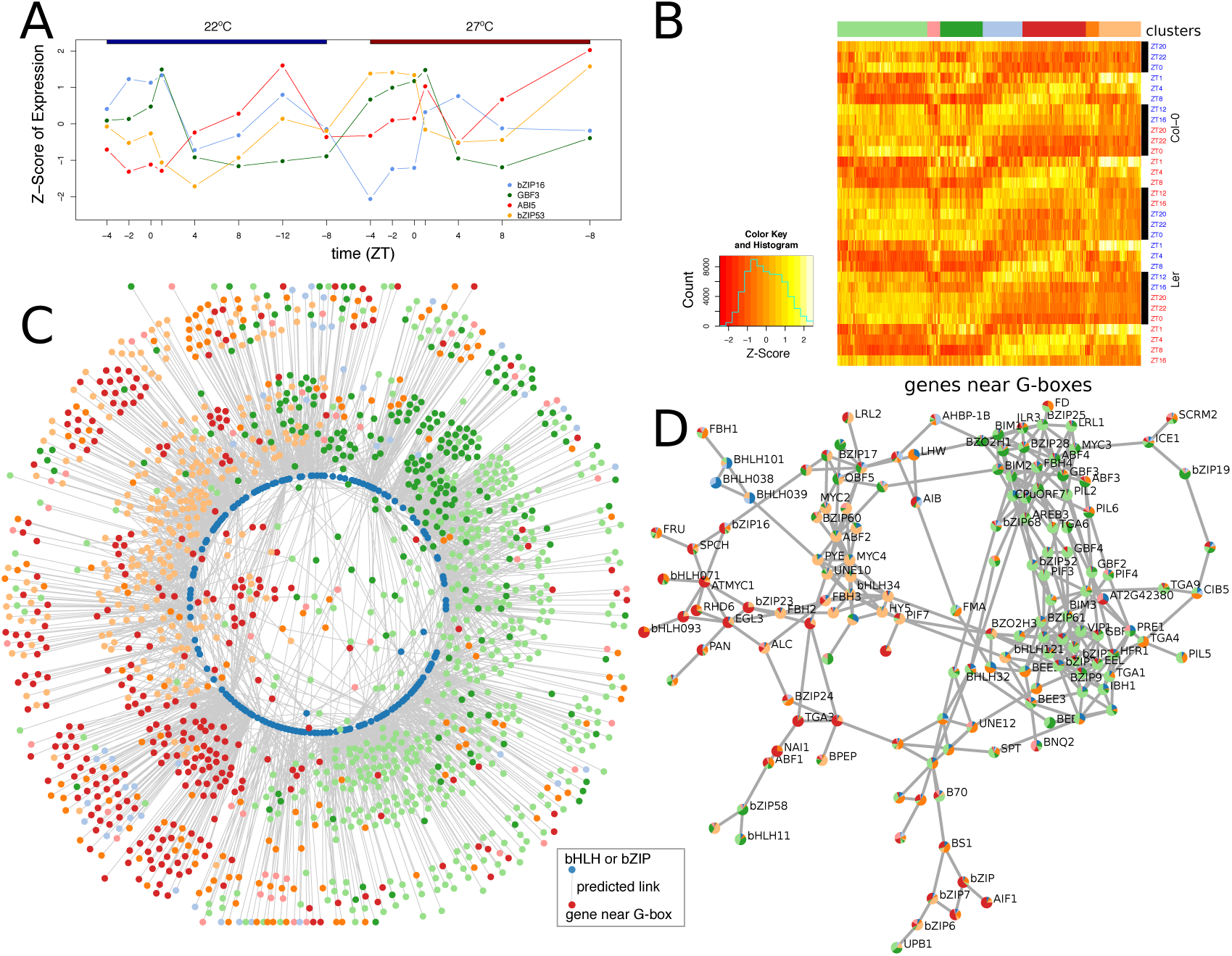
Gene expression pattern of genes near G-boxes used to infer regulatory network. (A) There are four bZIPs which bind to G-boxes in the “binding pattern 1” cluster of Figure 1A that are also expressed with TPM>1 in seedlings. They all have very different gene expression profiles. (B) Overall, most genes near G-boxes are expressed diurnally, and seven clusters were identified— the labeled colors of these clusters will be used consistently throughout the remainder of the paper. (C) This network depicts predicted regulatory links between bHLHs/bZIPs and genes near G-boxes are shown. Note that the network naturally reconstructs the circadian clock-- dawn genes, night genes, and day genes all cluster with their own kind. The colours are the same as in (B), except that TFs are shown in dark blue. (D) To condense the network, each TF was depicted as a pie chart indicating the time of day clusters of the genes it is predicted to regulate. The dawn cluster is particularly tight, and two distinct night clusters are also present.

We therefore constructed a gene regulatory network to determine the best set of bHLHs and bZIPs able to predict the gene expression of our set of genes downstream of perfect G-boxes. Compared to other co-expression networks, we would expect to see enrichment for direct TF-gene regulatory interactions, since we have prior knowledge that these TFs have the potential to bind and regulate this set of genes. The network was constructed based on a large set of RNA-seq time course data summarised in **Table S3**. These samples include *Arabidopsis* strains with loss of function mutations and over-expressors of genes involved in the circadian clock (*lux-4* and *elf3-1--* SRA:SUB1977196), temperature and red light signalling (*phyABCDE*-- SRA:PRJNA341458), histone remodelling (*dek3, DEK3-OX, hos1-3*, arp6—SRA:SUB2444423 and SUB2444427), and sugar processing (*ss4-1—* SRA:SUB2444427), as well as two wildtype strains (Col-0 and Ler—SRA:PRJNA341458 and SUB1977196). In most cases, gene expression data was collected in at least two different temperatures, usually 22^°^C and 27^°^C and sampled at eight or more time points throughout a 24-hour period. These RNA-seq datasets were selected primarily because they perturbed systems that were regulated by G-boxes, including temperature and light response and the circadian clock. In total, there were 229 RNA-seq samples utilised in the analysis. One RNA-seq time-course (for 8 time points of *pif4-101*, SRA:SUB 2444427, at 22^°^C and 27^°^C) is not used in this portion of the analysis, but is introduced later to help validate the network structure.

Clustering of the data revealed that almost all of the genes near G-boxes were expressed in a diurnal gene expression pattern (see **Fig 3B** for Col-0 and Ler only and **S5** for complete clustering). In particular, there are two large clusters of genes that were most highly expressed in the late night and dawn (light green and dark green) and another large cluster of genes that was expressed sharply an hour after dawn (tan)—this rapid induction of gene expression at dawn is referred to as the “morning peak” which has been conjectured to play a role in resetting the circadian clock in response to light (47).

From this extensive gene expression data, the network was inferred using an ensemble machine learning approach, by averaging the rank of edges predicted from a random forest approach (Genie3 (37)), a regression approach (Tigress (38)), and a mutual information approach (CLR (39)) (**Fig S6**). Such a network inference strategy was shown to be particularly strong in a crowd-sourced network inference challenge (40), and we found that all three methods consistently predicted a similar set of edges (**Fig S7**). The resulting gene expression network naturally reconstructs the circadian cycle when drawn with a force-directed layout and with the position of TF nodes (indicated by dark blue circles) restricted to a small inner circle (**Fig 3C**). In order to further probe the core structure of the network, we visualised the TF-TF interactions in the network, drawing each TF as a pie chart illustrating the proportion of its predicted targets fall into each cluster (**Fig 3D**). It is important to note that if two TFs were both predicted to target the same gene, they would have been ‘pulled’ together by the force-directed layout in Figure 3C, but not in Figure 3D, unless it was also predicted that one TF directly regulates the other TF. In such a depiction, the day-night cycle structure of the network is present, but less clearly visible. Instead, the most striking feature is that three strong clusters of nodes emerge—one tight group of dawn-expressed genes and two separate groups of genes that are expressed primarily at night. In contrast, the daytime genes appear more dispersed. Interestingly, the TFs that link the two clusters of night genes are PHYTOCHROME INTERACTING FACTORS (PIFs): PIF4, PIF3, PIF5, and PIF6. PIFs have emerged as key G-box binding TFs, since they integrate environmental signals to control morphogenesis in the late night (48). These results suggest that PIFs may have a key role in the structure of the plant regulatory network: linking the early night and late night/dawn transcriptional programs.

If there is a group of TFs that all have very similar expression patterns, then there is a high degree of ‘redundant’ information in that portion of the network. For example, since there is a substantial set of TFs that are part of the dawn cluster, these TFs convey ‘redundant’ information and there would be many possible choices of TFs that could be used to accurately predict the expression of genes expressed in the dawn peak. In order to quantify how much redundancy there is in the network, we calculate how easy it is to predict the gene expression pattern of each gene near a G-box, using the gene expression of a random subset of TFs as input (**Fig S8-A**). As expected, the most predictable gene expression profiles come from genes that are highly connected in the network (**Fig S8-B**) or that are part of dense subnetworks representing night or at dawn (**Fig S8-C**).

### The network can be used to predict gene expression patterns in a PHYTOCHROME INTERACTING FACTOR4 (PIF4) knockout

We next sought to determine whether our model could correctly predict gene expression patterns after a gene perturbation—i.e. whether the network is capable of predicting the gene expression arising from data that was not utilised when constructing the network. To test this, we evaluated the ability of the network to predict the gene expression pattern in a *pif4-101* time course at 22^°^C and 27^°^C, which was not used in the construction of the original network. For each gene near a G-box, we trained a random forest model that uses as input the gene expression pattern of TFs that it’s linked to in the network, again excluding the *pif4-101* data in the training. Finally, we used this random forest model to predict the gene expression in the *pif4-101* set and calculate its Pearson’s correlation coefficient. As a control, we train another model that uses the same number of TFs, but ones that are randomly sampled instead of those predicted by the network (**Fig S9**). If the network performed no better than a random network, we would expect these two models to predict the gene expression pattern of the *pif4-101* genes equally well. However, as shown in **Fig 4A**, the predicted network performs substantially better than random. Note that there is a large set of genes that do not have a substantial change in gene expression in the *pif4-101*—for these genes, it is easy to accurately predict the gene expression pattern in *pif4-101* using a random network. Critically, the larger the perturbation caused by the *pif4-101* mutant, the greater the improvement we see from the predicted network compared to the random network (**Fig S10**). If a gene has the same expression pattern in Col-0 and *pif4-101*, then even an over-fit random model would be likely to predict the expression of that gene accurately. However, if there were a large perturbation in gene expression, a network that captures something more fundamental about the co-expression of genes should perform better than random. Overall, this analysis indicates that the Ara-BOX-cis network can be used to predict gene expression, suggesting that its links/edges are biologically relevant.

**Figure 4:**
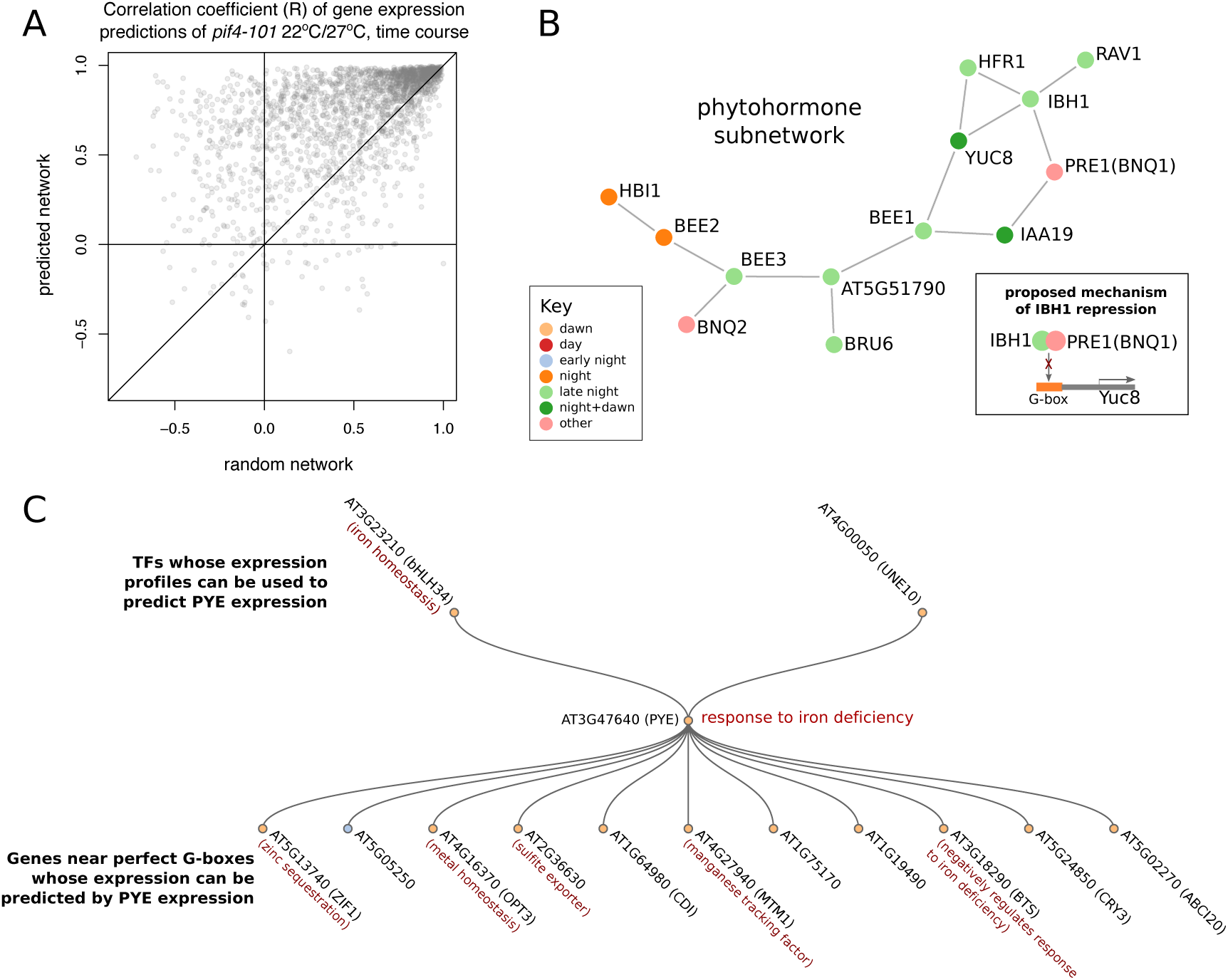
The Ara-BOX-sis regulatory network is predictive. (A) A time course of *pif4-101* RNA-seq experiments at 22^°^C and 27^°^C were not included in training the gene regulatory network. For each gene, a random forest regression model was trained to predict its gene expression based on the gene expression of the predicted regulatory links (or a random set of the same number of bZIP and bHLHs as a control). Both of these models were trained on the original RNA-seq data used to construct the regulatory network (i.e. not including *pif4-101*), but they were tested on the *pif4-101* RNA-seq dataset and the Pearson’s correlation coefficient was calculated. Points above the line indicate that the predicted network performed better than a random network at predicting new gene expression data. (B) The Ara-BOX-sis network includes a sub-network of phytohormone response genes with known biological interaction— see main text. Most notably, IBH1 is a bHLH that does not bind G-boxes, but that can heterodimerise with PRE1 and some BEE proteins to prevent them from binding to their targets, such as the G-box in the promoters of YUC8 and IAA19. (C) In addition, Ara-BOX-sis correctly identifies a known metal homeostasis subnetwork, centred on the PYE gene which is responsible for iron deficiency. Note that it directly interacts with BTS and many of its predicted targets change their gene expression in *pye*. It is important to note that no phytohormone or metal ions were directly influenced in the RNA-seq experiments; however, both of these processes are known to be temperature and circadian clock dependent.

### The network identifies metal homeostasis and phytohormone subnetworks

Another indication of the strength of our network inference approach is our ability to identify known sub-networks involved in systems that were not perturbed directly in any of the RNA-seq experiments.

One such subnetwork is involved in phytohormone response, which is particularly interesting since it lies at the threshold between the night and dawn clusters of the gene regulation network (**Fig 4B**). This is consistent with the hypothesis that hormone signalling helps regulate the dawn gene expression burst (47). Recently, it has been confirmed that LONG HYPOCOTYL IN FAR-RED (HFR1) directly regulates *YUCCA 8* (*YUC8*) (49). Moreover, BANQUO 1 (BNQ1) is a non-DNA binding bHLH that can antagonistically bind ILI1 BINDING BHLH 1 (IBH1) thereby preventing it from binding and regulating its downstream targets, a regulatory mechanism conserved in rice (50). Furthermore, recent work has identified that IAA19, BEE1/2/3, HBI1, IBH1, HFR1 are all part of the same gene expression network that link hormone, light response, and cell elongation (11).

Another interesting sub-network that was inferred by Ara-BOX-cis was the metal homeostasis network (**Fig 4C**), centred on the root iron homeostasis gene *POPEYE* (*PYE*), which appears to be regulated by the master iron homeostasis regulator bHLH34 (51). The predicted downstream targets of PYE include BTS, a tightly co-regulated protein that is also involved in iron homeostasis, and two direct targets of PYE-- *OBP3-RESPONSIVE GENE 1* (*ORG1*) and *ZINC TRANSPORTER 1 PRECURSOR* (*ZIF1*), as determined by a combination of microarray expression analysis and ChIP-on-chip (16).

Nevertheless, as this is a co-expression network, the links describe the correlations between G-box binding TF expression and the expression of genes downstream of G-boxes, and do not prove causation.

Although both phytohormone response and metal homeostasis are time-of-day dependent responses, neither system was specifically perturbed in any of the RNA-seq experiments used to infer the network. This indicates that the network has the potential to provide biologically relevant information to researchers who have an interest in particular G-box binders or genes near G-boxes.

### Ara-BOX-cis as a resource for plant biologists

To maximise the usefulness of Ara-BOX-cis to the community, we have made it available through a user-friendly web-browser (www.araboxcis.org). The Ara-BOX-cis network has three interactive layouts available for helping researchers generate testable hypothesis for the genes regulating a specific G-box or a set of genes regulated by G-boxes (**Fig S6**). Firstly, users can observe the entire network at once. In this view, they can drag around edges in order to more intuitively understand the make-up of the network, as well as identify motifs that are enriched in the predicted downstream targets of certain well-connected TFs and highlight genes based on the DNA sequences flanking the G-box. In the second layout, users can search for a particular gene via its TAIR ID and see the gene expression network centred on that gene—the gene’s description and the gene expression cluster of the parents and/or children of the node in the network are shown as in. Finally, a set of genes can be input by their TAIR IDs and the website will tally the number of times TFs or genes with G-boxes in their promoters appear one-step removed from any of these genes within the network.

## DISCUSSION

Understanding plant gene networks is complicated by the presence of large TF families (1), which is especially true in the case of bHLHs and bZIPs. There have been many instances where researchers have identified that a G-box was critical for the regulation of their gene of interest, but have been unable to pinpoint the exact TF or group of TFs that bind and regulate the target gene (22–26). In this paper, we present a set of resources to help researchers generate testable hypotheses in such a circumstance. Through analysis of TF-promoter interaction from DAP-seq data (35), and by observation of sequences flanking the G-box, it is possible to determine the *in vitro* binding specificity of bZIP homodimers. Then, by observing the position of a gene within the Ara-BOX-cis network it may be possible to predict which TFs are likely to regulate the gene of interest.

The network provided by Ara-BOX-cis was constructed using time-course RNA-seq samples that are relevant to biological processes related to G-box functions; it is considered the best practice for plant gene regulatory network reconstruction to use gene expression samples that are most likely to make large perturbations in the sub-network under study (52). It also uses network inference approaches that have been found to be most effective in side-by-side comparisons of network inference algorithms (40) and that has been previously been used to successfully reconstruct the root gene expression network in Arabidopsis and legumes (53, 54). Although it has fewer genes than other network approaches such as AraNET (55) and GeneMANIA (56), Ara-BOX-cis has the benefit of identifying possible links between TFs for which there is prior information suggesting a regulatory link, which is similar to the ‘weak prior’ approach suggested in Krouk et al, 2013 (57). All the TFs in this network either bind to G-boxes or heterodimerise with TFs that can do so, and all the other genes in the network have a perfect G-box within 500bp of their TSS. By limiting the predictions to this set, we enrich for direct regulatory associations.

We note that Ara-BOX-cis’s predicted edges may not necessarily represent direct regulatory interactions. For instance, there may be a bHLH and a gene near a G-box that are both regulated by the same non-G-box-binding TF. It is likely that Ara-BOX-cis will predict an edge between this bHLH and the gene near the G-box even if there is no direct interaction. This may be the case with the network centred on PIF5: the Evening Complex has been recently shown to regulate PIF5 and many of PIF5’s predicted targets (Ezer et al, 2017 submitted, and **Fig S11**). Nevertheless, the ability of Ara-BOX-cis to successfully predict the metal homeostasis and phytohormone sub-networks demonstrates that Ara-BOX-cis makes biologically relevant predictions.

There are many other conserved cis-regulatory boxes in *Arabidopsis*, such as C-boxes, W-boxes, and Heat Shock Elements, and a similar approach of analysing DAP-seq data and generating regulatory networks could be applied to these.

Understanding how plants regulate their genes is of profound importance. For instance, in the “Green Revolution” of agriculture that saved millions of lives from famine, one of the key achievements was the development of semi-dwarf wheat varieties. Later it was determined that this phenotype arose from a gene variant for the TF *GA REQUIRING 1* (58). This has sparked interest in the development of new biotechnologies that target TFs to further improve yields and also offer resilience to environmental fluctuations (59, 60). Understanding how transcription factor duplications affects plant gene regulation is critical if we hope to reverse engineer plant regulatory networks for agricultural applications.

## METHODS

### RNA-seq experiments and data processing

Data for Col-0, *elf3-1*, and *lux-4* is discussed in (Ezer et al, 2017, submitted). Data for Ler, phyABCDE, YHB is available from Jung et al 2016 (14). The RNA-seq experiments on the *ss4-1, hos1-3*, and *pif4-101* lines are first presented here. These lines were all described previously: *ss4-1* in (27), *hos1-3* in (28), and *pif4-101* in (29). It is a T-DNA insertion mutant in a Col-0 background. Seeds were sown on ½ X Murashige and Skoog-agar (MS-agar) plates at pH 5.7 without sucrose, stratified at 4°C in the dark for 3 days and germinated for 24 h at 22°C. Plants were then put into Conviron PGC20 reach-in growth cabinets under 170 μmol m^2^ s^-1^ white light at either 22°C or 27°C short days. Plants were then sampled at various points across a 24 h timecourse over the diurnal cycle: ZT = 0, 1, 4, 8, 12, 16, 20 and 22 h. RNA was extracted using the MagMax-96 Total RNA extraction kit (Ambion, AM1830). RNA quality and integrity was assessed on the Agilent 2200 TapeStation. Library preparation was performed using 1 μg of high integrity total RNA (RIN>8) using the TruSeq RNA Library Preparation Kit v2 (Illumina, RS-122-2101 and RS-122-2001), following the manufacturer’s instruction. The libraries were sequenced on a HiSeq2000 using paired-end sequencing of 100 bp in length at the Beijing Genomics Institute (BGI) sequencing center.

The same pipeline was used to map these sequences as described in previously(14). To analyse the sequence reads: First, adapters were trimmed with Trimmomatic-0.32(30). Then, Tophat(31) was used to map to the TAIR10 annotated genome, duplicates were removed and the read counts were normalised by genome-wide coverage. Raw counts were determined by HTseq-count(32), and cufflinks was used to calculate Fragments Per Kilobase Million (FPKM), which was then converted into Transcripts Per Million (TPM). Note that this protocol for RNA-seq and data processing is the same as the one described in Jung et al, 2016 (14) and Ezer et al, 2017, submitted.

### MNase-seq experiments and data processing

Col-0 WT Arabidopsis seedlings were grown on the ½ MS plates in short day conditions 8hr light/16hr dark at 17^°^C. Ten-day old seedlings were harvested 8hr after dawn. Plant material was immediately cross-linked using 1% formaldehyde under vacuum. Nuclei were purified as previously described (33). Chromatin, extracted from 1 g of materials, was resuspended in MNase digestion buffer (20mM Tris-HCl [pH8], 50mM NaCl, 1mM DTT, 0.5% NP-40, 1mM CaCl2, 0.5mM phenylmethylsulfonyl fluoride (PMSF) and 1X protease inhibitor cocktail [Roche]), and digested with 0.4U/ml of micrococcal nuclease (MNase, Sigma, N3755) for 15 min. MNase digestion was performed to obtain a mono-nucleosome resolution as MNase preferentially digests nucleosome-free DNA and the linker regions, whereas sequences bound by nucleosomes are relatively protected from the digestion (34).

The efficiency of MNase was assessed by separating samples on the agarose gel in prior to in-house library preparation, using the TruSeq ChIP sample preparation kit (Illumina, IP-202-1012). The libraries were sequenced using paired-end 75bp on NextSeq500 on site.

### Selection of genes

The list of bZIPs and bHLHs came from TAIR10 annotations (https://www.arabidopsis.org/browse/genefamily/bZIP.jsp and https://www.arabidopsis.org/browse/genefamily/bHLH.jsp).

For the identification of genes that are likely regulated by G-boxes in seedlings, genes with transcripts per million (TPM)>1 in at least half of our primary set of RNA-seq time-courses: Col-0, Ler, *elf3-1, lux-4*, and *phyABCDE* were initially selected. The 500bp DNA sequences upstream of their TSS were extracted from TAIR10, and the sequence CACGTG was searched for using an R script available on github.

### DAP-seq data

The DAP-seq data were extracted from neomorph.salk.edu/PlantCistromeDB (35), and additional bHLH datasets (bHLH18, bHLH34, bHLH69, bHLH74, bHLH77, bHH104, BIM3, PIF7) were taken directly from GEO (**GSE60141**). In particular, the bed files listing significant TF binding peaks (**Table S2**) were searched using bedtools intersect and the heatmap was drawn in R.

### Random Forest and DNA Shape

In all cases, R’s randomforest package was used with the number of trees set to 1000. These scripts are available on Github, and the experimental designs for each instance random forest was used is drawn in the Supplemental Figures referenced in the text. Note that the DNA shape was calculated using the DNAShapeR package (36).

### Ara-BOX-cis

Ara-BOX-cis utilised Genie3 (37),Tigress (38) and CLR (39). The ranks of each edge were averaged—a procedure suggested in (40). This was done in R, but note that all ranks that were over 20,000 were set to have a value of 20,000. The threshold for average rank used for drawing the figures in **Fig 3C** is 5,000 and the threshold for average rank in **Fig 3D** is 10,000. The network was visualised and the website was constructed using d3.js (https://d3js.org/), in particular the two-way tree example was used as a template for one panel of the website (http://bl.ocks.org/kanesee/5d6c48bffd4ea31201fb).

## AVAILABILITY

The Ara-BOX-cis network is available on araboxcis.org. All other code used to analyse this data is available on github (https://github.com/ezer/GboxRscripts). Data structures/tables to assist in running the code efficiently are available in https://github.com/ezer/Gboxdata. The contents of the Data folder from the Gboxdata repository can be added to the home folder of the GboxRscripts repository before running the code.

## ACCESSION NUMBERS

All raw RNA-seq data and MNase-seq data will be made publicly available in SRA: (SUB2444423, SUB2445017, SUB1977196, PRJNA341458, SUB2444427).

## ACKNOWLEDGEMENT

We thank members of the Wigge laboratory for feedback and discussions.

## REFERENCES

1. Shiu, S.-H., Shih, M.-C. and Li, W.-H. (2005) Transcription factor families have much higher expansion rates in plants than in animals. Plant Physiol., 139, 18–26.

2. Charoensawan, V., Wilson, D. and Teichmann, S.A. (2010) Genomic repertoires of DNA-binding transcription factors across the tree of life. Nucleic Acids Res., 38, 7364–7377.

3. Gordân, R., Shen, N., Dror, I., Zhou, T., Horton, J., Rohs, R. and Bulyk, M.L. (2013) Genomic regions flanking E-box binding sites influence DNA binding specificity of bHLH transcription factors through DNA shape. Cell Rep., 3, 1093–1104.

4. Nuruzzaman, M., Sharoni, A.M. and Kikuchi, S. (2013) Roles of NAC transcription factors in the regulation of biotic and abiotic stress responses in plants. Front. Microbiol., 4, 248.

5. Altman, B.J., Hsieh, A.L., Sengupta, A., Krishnanaiah, S.Y., Stine, Z.E., Walton, Z.E., Gouw, A.M., Venkataraman, A., Li, B., Goraksha-Hicks, P., et al. (2015) MYC Disrupts the Circadian Clock and Metabolism in Cancer Cells. Cell Metab., 22, 1009–1019.

6. Rensing, S.A. (2014) Gene duplication as a driver of plant morphogenetic evolution. Curr. Opin. Plant Biol., 17, 43–48.

7. Carretero-Paulet, L., Galstyan, A., Roig-Villanova, I., Martínez-García, J.F., Bilbao-Castro, J.R. and Robertson, D.L. (2010) Genome-wide classification and evolutionary analysis of the bHLH family of transcription factors in Arabidopsis, poplar, rice, moss, and algae. Plant Physiol., 153, 1398–1412.

8. Heim, M.A., Jakoby, M., Werber, M., Martin, C., Weisshaar, B. and Bailey, P.C. (2003) The basic helix-loop-helix transcription factor family in plants: a genome-wide study of protein structure and functional diversity. Mol. Biol. Evol., 20, 735–747.

9. Corrêa, L.G.G., Riaño-Pachón, D.M., Schrago, C.G., dos Santos, R.V., Mueller-Roeber, B. and Vincentz, M. (2008) The role of bZIP transcription factors in green plant evolution: adaptive features emerging from four founder genes. PLoS One, 3, e2944.

10. Hao, Y., Oh, E., Choi, G., Liang, Z. and Wang, Z.-Y. (2012) Interactions between HLH and bHLH factors modulate light-regulated plant development. Mol. Plant, 5, 688–697.

11. Oh, E., Zhu, J.-Y., Bai, M.-Y., Arenhart, R.A., Sun, Y. and Wang, Z.-Y. (2014) Cell elongation is regulated through a central circuit of interacting transcription factors in the Arabidopsis hypocotyl. Elife, 3.

12. Toledo-Ortiz, G., Johansson, H., Lee, K.P., Bou-Torrent, J., Stewart, K., Steel, G., Rodríguez-Concepción, M. and Halliday, K.J. (2014) The HY5-PIF regulatory module coordinates light and temperature control of photosynthetic gene transcription. PLoS Genet., 10, e1004416.

13. Choi, H. and Oh, E. (2016) PIF4 Integrates Multiple Environmental and Hormonal Signals for Plant Growth Regulation in Arabidopsis. Mol. Cells, 39, 587–593.

14. Jung, J.-H., Domijan, M., Klose, C., Biswas, S., Ezer, D., Gao, M., Khattak, A.K., Box, M.S., Charoensawan, V., Cortijo, S., et al. (2016) Phytochromes function as thermosensors in Arabidopsis. Science, 354, 886–889.

15. Alves, M.S., Dadalto, S.P., Gonçalves, A.B., De Souza, G.B., Barros, V.A. and Fietto, L.G. (2013) Plant bZIP transcription factors responsive to pathogens: a review. Int. J. Mol. Sci., 14, 7815–7828.

16. Long, T.A., Tsukagoshi, H., Busch, W., Lahner, B., Salt, D.E. and Benfey, P.N. (2010) The bHLH transcription factor POPEYE regulates response to iron deficiency in Arabidopsis roots. Plant Cell, 22, 2219–2236.

17. Nakashima, K., Yamaguchi-Shinozaki, K. and Shinozaki, K. (2014) The transcriptional regulatory network in the drought response and its crosstalk in abiotic stress responses including drought, cold, and heat. Front. Plant Sci., 5, 170.

18. Gangappa, S.N., Maurya, J.P., Yadav, V. and Chattopadhyay, S. (2013) The regulation of the Z-and G-box containing promoters by light signaling components, SPA1 and MYC2, in Arabidopsis. PLoS One, 8, e62194.

19. Zhou, T., Shen, N., Yang, L., Abe, N., Horton, J., Mann, R.S., Bussemaker, H.J., Gordân, R. and Rohs, R. (2015) Quantitative modeling of transcription factor binding specificities using DNA shape. Proc. Natl. Acad. Sci. U. S. A., 112, 4654–4659.

20. Swarnalatha, M., Singh, A.K. and Kumar, V. (2012) The epigenetic control of E-box and Myc-dependent chromatin modifications regulate the licensing of lamin B2 origin during cell cycle. Nucleic Acids Res., 40, 9021–9035.

21. Fernandez, P.C., Frank, S.R., Wang, L., Schroeder, M., Liu, S., Greene, J., Cocito, A. and Amati, B. (2003) Genomic targets of the human c-Myc protein. Genes Dev., 17, 1115–1129.

22. Liu, T.L., Newton, L., Liu, M.-J., Shiu, S.-H. and Farré, E.M. (2016) A G-Box-Like Motif Is Necessary for Transcriptional Regulation by Circadian Pseudo-Response Regulators in Arabidopsis. Plant Physiol., 170, 528–539.

23. Liu, L., Xu, W., Hu, X., Liu, H. and Lin, Y. (2016) W-box and G-box elements play important roles in early senescence of rice flag leaf. Sci. Rep., 6, 20881.

24. Kobayashi, K., Obayashi, T. and Masuda, T. (2012) Role of the G-box element in regulation of chlorophyll biosynthesis in Arabidopsis roots. Plant Signal. Behav., 7, 922–926.

25. Kim, K.N. and Guiltinan, M.J. (1999) Identification of cis-acting elements important for expression of the starch-branching enzyme I gene in maize endosperm. Plant Physiol., 121, 225–236.

26. Hwang, Y.S., Karrer, E.E., Thomas, B.R., Chen, L. and Rodriguez, R.L. (1998) Three cis-elements required for rice alpha-amylase Amy3D expression during sugar starvation. Plant Mol. Biol., 36, 331–341.

27. Roldán, I., Wattebled, F., Mercedes Lucas, M., Delvallé, D., Planchot, V., Jiménez, S., Pérez, R., Ball, S., D’Hulst, C. and Mérida, Á. (2007) The phenotype of soluble starch synthase IV defective mutants of Arabidopsis thaliana suggests a novel function of elongation enzymes in the control of starch granule formation. Plant J., 49, 492–504.

28. Jung, J.-H.J.-H., Park, J.-H., Lee, S., To, T.K., Kim, J.-M.J.-M., Seki, M. and Park, C.-M.C.-M. (2013) The Cold Signaling Attenuator HIGH EXPRESSION OF OSMOTICALLY RESPONSIVE GENE1 Activates FLOWERING LOCUS C Transcription via Chromatin Remodeling under Short-Term Cold Stress in Arabidopsis. Plant Cell, 25, 4378–4390.

29. Koini, M.A., Alvey, L., Allen, T., Tilley, C.A., Harberd, N.P., Whitelam, G.C. and Franklin, K.A. (2009) High Temperature-Mediated Adaptations in Plant Architecture Require the bHLH Transcription Factor PIF4. Curr. Biol., 19, 408–413.

30. Bolger, A.M., Lohse, M. and Usadel, B. (2014) Trimmomatic: a flexible trimmer for Illumina sequence data. Bioinformatics, 30, 2114–20.

31. Trapnell, C., Pachter, L. and Salzberg, S.L. (2009) TopHat: discovering splice junctions with RNA-Seq. Bioinformatics, 25, 1105–11.

32. Anders, S., Pyl, P.T. and Huber, W. (2015) HTSeq--a Python framework to work with high-throughput sequencing data. Bioinformatics, 31, 166–9.

33. Folta, K.M. and Kaufman, L.S. (2006) Isolation of Arabidopsis nuclei and measurement of gene transcription rates using nuclear run-on assays. Nat. Protoc., 1, 3094–3100.

34. Petesch, S.J. and Lis, J.T. (2008) Rapid, Transcription-Independent Loss of Nucleosomes over a Large Chromatin Domain at Hsp70 Loci. Cell, 134, 74–84.

35. O’Malley, R.C., Huang, S.-S.C., Song, L., Lewsey, M.G., Bartlett, A., Nery, J.R., Galli, M., Gallavotti, A. and Ecker, J.R. (2016) Cistrome and Epicistrome Features Shape the Regulatory DNA Landscape. Cell, 165, 1280–1292.

36. Chiu, T.-P., Comoglio, F., Zhou, T., Yang, L., Paro, R. and Rohs, R. (2016) DNAshapeR: an R/Bioconductor package for DNA shape prediction and feature encoding. Bioinformatics, 32, 1211–1213.

37. Huynh-Thu, V.A., Irrthum, A., Wehenkel, L. and Geurts, P. (2010) Inferring regulatory networks from expression data using tree-based methods. PLoS One, 5.

38. Haury, A.-C., Mordelet, F., Vera-Licona, P. and Vert, J.-P. (2012) TIGRESS: Trustful Inference of Gene REgulation using Stability Selection. BMC Syst. Biol., 6, 145.

39. Madar, A., Greenfield, A., Vanden-Eijnden, E. and Bonneau, R. (2010) DREAM3: network inference using dynamic context likelihood of relatedness and the inferelator. PLoS One, 5, e9803.

40. Marbach, D., Costello, J.C., Küffner, R., Vega, N.M., Prill, R.J., Camacho, D.M., Allison, K.R., Consortium, D., Kellis, M., Collins, J.J., et al. (2012) Wisdom of crowds for robust gene network inference. Nat. Methods, 9, 796–804.

41. He, J.-X., Gendron, J.M., Sun, Y., Gampala, S.S.L., Gendron, N., Sun, C.Q. and Wang, Z.-Y. (2005) BZR1 is a transcriptional repressor with dual roles in brassinosteroid homeostasis and growth responses. Science, 307, 1634–1638.

42. Zentner, G.E. and Henikoff, S. (2012) Surveying the epigenomic landscape, one base at a time. Genome Biol., 13, 250.

43. Henikoff, J.G., Belsky, J.A., Krassovsky, K., MacAlpine, D.M. and Henikoff, S. (2011) Epigenome characterization at single base-pair resolution. Proc. Natl. Acad. Sci. U. S. A., 108, 18318–18323.

44. Guiltinan, M.J., Marcotte, W.R. and Quatrano, R.S. (1990) A plant leucine zipper protein that recognizes an abscisic acid response element. Science, 250, 267–271.

45. Williams, M.E., Foster, R. and Chua, N.H. (1992) Sequences flanking the hexameric G-box core CACGTG affect the specificity of protein binding. Plant Cell, 4, 485–496.

46. Para, A., Li, Y., Marshall-Colón, A., Varala, K., Francoeur, N.J., Moran, T.M., Edwards, M.B., Hackley, C., Bargmann, B.O.R., Birnbaum, K.D., et al. (2014) Hit-and-run transcriptional control by bZIP1 mediates rapid nutrient signaling in Arabidopsis. Proc. Natl. Acad. Sci. U. S. A., 111, 1–6.

47. Michael, T.P., Breton, G., Hazen, S.P., Priest, H., Mockler, T.C., Kay, S.A. and Chory, J. (2008) A morning-specific phytohormone gene expression program underlying rhythmic plant growth. PLoS Biol., 6, e225.

48. Leivar, P. and Monte, E. (2014) PIFs: systems integrators in plant development. Plant Cell, 26, 56–78.

49. Hayes, S., Sharma, A., Fraser, D.P., Trevisan, M., Cragg-Barber, C.K., Tavridou, E., Fankhauser, C., Jenkins, G.I. and Franklin, K.A. (2016) UV-B Perceived by the UVR8 Photoreceptor Inhibits Plant Thermomorphogenesis. Curr. Biol., 10.1016/j.cub.2016.11.004.

50. Zhang, L.-Y., Bai, M.-Y., Wu, J., Zhu, J.-Y., Wang, H., Zhang, Z., Wang, W., Sun, Y., Zhao, J., Sun, X., et al. (2009) Antagonistic HLH/bHLH transcription factors mediate brassinosteroid regulation of cell elongation and plant development in rice and Arabidopsis. Plant Cell, 21, 3767–3780.

51. Li, X., Zhang, H., Ai, Q., Liang, G. and Yu, D. (2016) Two bHLH Transcription Factors, bHLH34 and bHLH104, Regulate Iron Homeostasis in Arabidopsis thaliana. Plant Physiol., 170, 528–539.

52. Gaudinier, A. and Brady, S.M. (2016) Mapping Transcriptional Networks in Plants: Data-Driven Discovery of Novel Biological Mechanisms. Annu. Rev. Plant Biol., 67, 575–594.

53. Taylor-Teeples, M., Lin, L., de Lucas, M., Turco, G., Toal, T.W., Gaudinier, A., Young, N.F., Trabucco, G.M., Veling, M.T., Lamothe, R., et al. (2015) An Arabidopsis gene regulatory network for secondary cell wall synthesis. Nature, 517, 571–575.

54. Wang, M., Verdier, J., Benedito, V.A., Tang, Y., Murray, J.D., Ge, Y., Becker, J.D., Carvalho, H., Rogers, C., Udvardi, M., et al. (2013) LegumeGRN: a gene regulatory network prediction server for functional and comparative studies. PLoS One, 8, e67434.

55. Lee, T., Yang, S., Kim, E., Ko, Y., Hwang, S., Shin, J., Shim, J.E., Shim, H., Kim, H., Kim, C., et al. (2015) AraNet v2: an improved database of co-functional gene networks for the study of Arabidopsis thaliana and 27 other nonmodel plant species. Nucleic Acids Res., 43, D996–1002.

56. Mostafavi, S., Ray, D., Warde-Farley, D., Grouios, C. and Morris, Q. (2008) GeneMANIA: a real-time multiple association network integration algorithm for predicting gene function. Genome Biol., 9 Suppl 1, S4.

57. Krouk, G., Lingeman, J., Colon, A., Coruzzi, G. and Shasha, D. (2013) Gene regulatory networks in plants: learning causality from time and perturbation. Genome Biol., 14, 123.

58. Peng, J., Richards, D.E., Hartley, N.M., Murphy, G.P., Devos, K.M., Flintham, J.E., Beales, J., Fish, L.J., Worland, A.J., Pelica, F., et al. (1999) ‘Green revolution’ genes encode mutant gibberellin response modulators. Nature, 400, 256–261.

59. Century, K., Reuber, T.L. and Ratcliffe, O.J. (2008) Regulating the regulators: the future prospects for transcription-factor-based agricultural biotechnology products. Plant Physiol., 147, 20–29.

60. Grotewold, E. (2008) Transcription factors for predictive plant metabolic engineering: are we there yet? Curr. Opin. Biotechnol., 19, 138–144.

